# Friend or Foe: Ambrosia Beetle response to volatiles of common threats in their fungus gardens

**DOI:** 10.1101/2022.12.23.521835

**Authors:** Janina M.C. Diehl, Denicia Kassie, Peter H.W. Biedermann

**Author notes:** <u>Authors for correspondence:</u> Janina M. C. Diehl, Peter H. W. Biedermann.

## Abstract

Fungus farming insects encounter multiple microbial threats in their cultivar gardens. They can affect both the nutritional cultivar and the insect’s health. In this study, we explored the potential of ambrosia beetles and their larvae to detect the presence of ubiquitous weed or entomopathogenic fungi. The ability to recognize a threat offers individuals a chance to react. Our study organism, the fruit-tree pinhole borer, *Xyleborinus saxesenii*, is associated with two mutualistic fungi, *Dryadomyces sulphureus* (*Raffaelea sulphurea)* and *Raffaelea canadensis*. Both symbionts were tested in combinations with two common fungus-garden weeds (*Aspergillus* sp. and *Penicillium commune*) and the entomopathogen *Beauveria bassiana* in two-choice experiments. Behavioural repellence was found in many, but not all combinations. Larvae and adult females showed an opposite reaction towards the entomopathogen, whereas for *Aspergillus* sp., neither provoked repellence nor attraction of larvae and adult females, if *R. canadensis* was used as lure. Our results validate a response of both larvae and adult ambrosia beetles towards other fungal volatiles. Their decision to confront a potential threat or preferably to avoid it could be subject to a more complex context.

## Introduction

Farming is a prominent behaviour found in several groups in the animal kingdom. Among the most prominent farmers, apart from humans, are fungus-cultivating insects (Mueller et al. 2005; Mayer et al. 2018; Biedermann and Vega 2020). A shared challenge of humans, as well as insects, is the ubiquitous threat of weeds and pathogens for the stable cultivation of crops (Schultz et al. 2022). Fungus farming insects encounter multiple threats of microbes in their cultivar gardens. These threats can be either a danger to the nutritional cultivar (e.g. competing “weeds” or pathogens (Schultz 2022)) or directly affect the health of the insects (e.g. entomopathogens (Kushiyev et al. 2018)). Ants and termites are known for their ability to detect and react to these intruders (Rosengaus et al. 1999; Griffiths and Hughes 2010; Katariya et al. 2017; Goes et al. 2020), but little is known about a third big group of fungus-farming, wood-boring weevils, the so called bark and ambrosia beetles (Curculionidae: Platypodinae and Scolytinae).

Bark and ambrosia beetles bore tunnels in phloem or xylem of trees, which they inoculate with mutualistic fungi, their sole food source (Harrington 2005; Kirkendall et al. 2015). Xylem-boring ambrosia beetles are obligately dependent on these fungi, while most bark beetles (except a few exceptions, e.g. *Dendroctonus* spp.) are facultatively associated with fungi (Six and Klepzig 2021). Whether all bark and ambrosia beetles are able to detect and react to fungal weeds is doubtful, but recent experiments on the ambrosia beetle, *Xyleborinus saxesenii* (Ratzeburg), revealed that this species can detect and selectively remove an *Aspergillus* fungal pathogen (Nuotclá et al. 2019). Furthermore, beetles can actively promote their nutritional mutualists over fungal weeds (Diehl et al. 2022). So far, it is unknown, however, how *X. saxesenii* and other farming beetles detect the presence of pathogens within nests. Volatiles of the weeds itself or changing volatile profiles of the nutritional fungi may be cues or signals, respectively. Attraction of bark and ambrosia beetles to fungal volatiles is still very sparsely documented. By now we know, that bark and ambrosia beetles can detect olfactory cues of their mutualists and are attracted to them, whereas non-associated, weedy fungi provided no sign of attraction (Hanula et al. 2008; Hulcr et al. 2011; Luna et al. 2014; Kandasamy et al. 2016; Kandasamy et al. 2019). Therefore, it is unclear if the beetles can detect the weedy fungi if lab studies just showed that they are not attracted (e.g. Hulcr et al. 2011). This requires the testing of weed fungi in combination with an attractive lure, which would show if its attraction is reduced in the presence of a fungal weed. Here, we tested whether both life stages (larvae and adult beetles) of the fruit-tree pinhole borer, *X. saxesenii*, can detect the presence of two fungus-garden competitors (*Aspergillus* sp. and *Penicillium commune*) and one entomopathogen (*Beauveria bassiana*), if presented together with its *Raffaelea* fungal mutualists, which attraction to adult beetles has been shown previously (Hulcr et al. 2011). Repellence of beetles would be expected in all three cases.

## Material/Methods

### Beetle breeding

In January 2022, nests were established in tubes filled with artificial beech medium following the standard protocol (Biedermann et al, 2009). Previously sib-mated *X. saxesenii* females out of our laboratory stock (origin: EtOH-baited trap-collected dispersers from Steinbachtal/Wuerzburg, Bavaria, June-July 2019) were surface-sterilized with 70% EtOH, followed by washing with distilled water and drying on a paper towel. Females were afterwards individually introduced into the sterile rearing medium and kept under standard conditions in the climate chamber (25°C, 24h darkness, 60-70% humidity) to establish a new generation of nests.

After some weeks, adult beetles and larvae were extracted for the experimental tests in February and March 2022 (*N* = 22 nests). The beetles used in the experiment were intended to be in the same period of development (approx. 28 days after nest foundation). In this developmental phase adult individuals had not dispersed out of the nest yet, but instead are engaged in fungus farming and brood care in the maternal nest (Nuotclà et al. 2019). Therefore, it is expected that they would be sensitive for the presence of fungal pathogens.

### Fungus Cultivation

Strains of the different mutualistic and antagonistic fungi used in this experiment were obtained from our laboratory’s stock of the Chair of Forest Entomology and Protection (Univ. Freiburg, Germany), stored on agar slants within tubes at 5°C and previously grown on yeast extract malt agar (YEMA; Carl Roth^®^) with streptomycin. We cultivated the different fungi under a sterile bench, by transferring a small sample of the fungi from the stock tubes with a sterilized scalpel onto the middle of a petri dishes with YEMA medium. Cultures were sealed with parafilm and incubated at 25°C.

Over time, several cultures of each *Dryadomyces sulphureus* (aka *Raffaelea sulphurea*; mutualist, GenBank Accession: MT880108.1), *R. canadensis* (mutualist, GenBank Accession: MT880109.1), *Penicillium commune* (antagonist, strain GenBank Accession: MT252032.1), *Aspergillus* sp. (antagonist, GenBank Accession: MT252035.1) and *Beauveria bassiana* (entomopathogen, GenBank Accession: MT159433.1) were weekly re-cultivated on new media to secure the permanent supply of fungal plugs for the experimental execution.

### Two-choice experiment with *X. saxesenii* adults

We performed 210 two-choice experiment runs with modified petri dishes for adult ambrosia beetles. Here, the surface of the dishes was roughened using sandpaper to allow unhindered walking. Furthermore, we perforated the petri dish three times – in equal distance to the center and each other – with a Dremel 2001 (Dremel Deutschland) to fit 1.5 ml Eppendorf reaction tubes (Fig. 1A/B). These tubes were fixed with glue and served both as cavities for fungal test plugs and collection vials for the choosing beetles. Since this was a two-choice set-up, only two tubes were punctured with a small hole, to allow beetles to detect volatiles and access the fungi. The third tube was used only as a stand and for stability of the construction. Before each test run, modified petri dishes were washed in EtOH and distilled water and placed in the sterile bench under a UV lamp (254 nm wave length) for 30 min.

**Figure 1:**
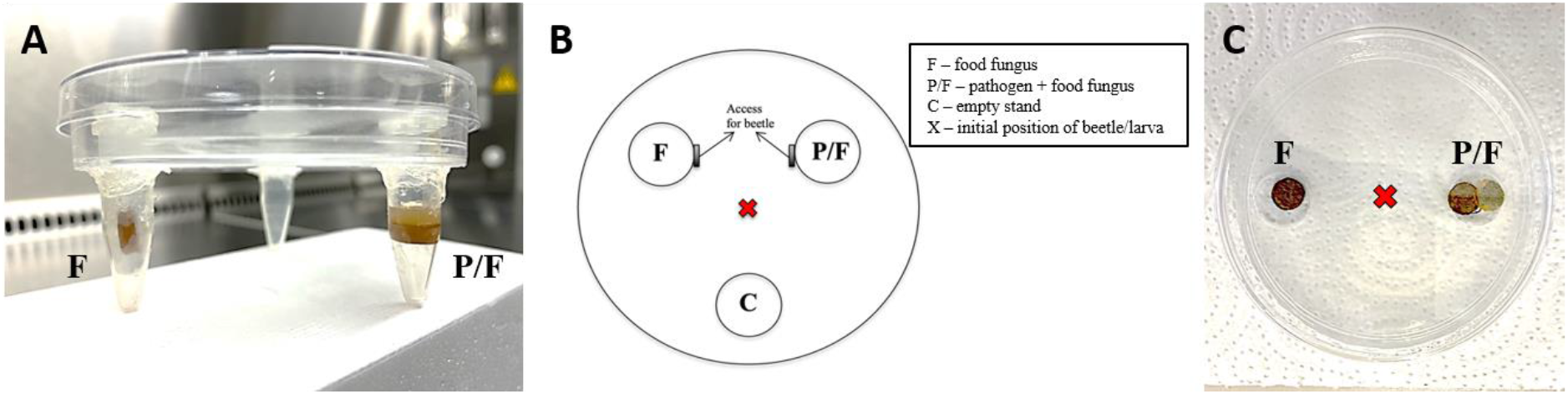
Experimental set-up of two-choice tests for *Xyleborinus saxesenii* females (A/B) and larvae (C). Petri dishes with a roughened surface were provided with three 1.5 ml Eppendorf tubes, two tubes included either one 5 mm plug of a food fungus or two plugs with each food fungus and pathogen to test female choice (A). Schematic representation of the experimental set-up for female choice tests (B). Petri dishes filled with solid agar-agar medium served as basis for the larval choice test. Here, two pits were punched out to place either one 5 mm plug of a food fungus or two plugs with each food fungus and pathogen (C).

We measured repellence of adult females to volatiles of the fungal pathogens by providing the test tubes either with a sole 5 mm diameter agar plug (cork borer) of one of the mutualistic food fungi (*D. sulphureus* or *R. canadensis*) (= positive control) or a food fungus in combination with one of the other antagonistic or entomopathogenic fungi (*P. commune, Aspergillus* sp. or *B. bassiana*) (= repellence treatment). In total we tested six combinations with each 35 replications. Combinations applied were: i) *D. sulphureus* and *P. commune*, ii) *R. canadensis* and *P. commune*, iii) *D. sulphureus* and *Aspergillus* sp., iv) *R. canadensis* and *Aspergillus* sp. v) *D. sulphureus* and *B. bassiana*, vi) *R. canadensis* and *B. bassiana*. Test individuals (females) were directly extracted from nests and placed after a surface sterilization in the center of the test arena. The choice-behaviour of focal beetles was observed for 30 min in 5-minute intervals and a final observation after 24 hours in our climate chamber under red light (switched on only during observations). Orientation of the arenas was changed regularly. Finally, we recorded, if beetles made a choice (‘choice’ by entering a tube or ‘no choice’) and if so, which side they chose (‘food’ or ‘food-pathogen’).

### Two-choice experiment on *X. saxesenii* larvae

Since larvae are slower moving and not protected by a chitinous carapace, they are more at risk of drying out during the experiment. Therefore, we used another set-up to test them (similar to Luna et al. 2014). Each arena was built of a petri dish with a bottom layer (1 cm) of solid agar-agar (14 g/800 ml, Carl Roth^®^) and two pits punched out with an eleven millimeter (diameter) cork borer on opposite sides. We provided each of these pits with either a plug (5 mm diameter) of the food fungus alone (= positive control) or a combination of food fungus and antagonist/entomopathogen (= repellence treatment; see above) (Fig. 1C). However, the secondary food fungus (*R. canadensis*) was not used in this series of tests, which resulted in only three combinations to be tested (*P. commune, Aspergillus* sp. and *B. bassiana*). We only used *D. sulphureus*, since larvae are more attracted to it, probably because it is their primary food source (Lehenberger; unpubl. data). Each combination was repeated 40 times with each a new petri dish (*N* = 120). To avoid arenas to dry out during tests, they were prepared three days ahead. We placed a single larva into the center of the plate after surface sterilization and recorded the position of it after 24 hours. If the larva was within a 1cm proximity to the pit, this was recorded as its final choice. Again, we distinguished between ‘choice’ and ‘no choice’, as well as ‘food’ and ‘food-pathogen’ for the final analysis.

### Statistics

All statistical analyses and visualizations were performed in RStudio (Version 1.4.1106) with R version 4.0.5 (R Core Team 2021) using the ‘ggplot2’ package (Wickham 2016) for the graphical output.

We ran a series of *chi-square* tests to examine if intended choices were made by the individuals per tested combination and life stage. To check if the sample size is good enough to perform appropriate tests, we looked at the expected values. Since all tested combinations had expected values the cells of the contingency table higher than 5, we were able to apply *chi-square* tests.

## Results

The food fungi were attractive in all combinations because all tested life-stages showed a statistically significant decision for a choice, except larvae in the combination with *D. sulphureus* and *B. bassiana*, which was almost significant (p = 0.114; Tab. 1). Behavioural repellence was found in many, but not all combinations. First, there was neither repellence nor attraction found in the reaction of larvae towards *Aspergillus* sp. and in adult females, if *R. canadensis* was used as lure (Tab. 1, Fig. 2). Second and most surprisingly, we found attraction of *B. bassiana*, in both combinations when either presented with *D. sulphureus* or *R. canadensis*. By contrast, larvae are repelled by this entomopathogenous fungus.

**Table 1:** Outcome of two-choice tests between food fungus and food fungus with pathogen for adult females and larvae. Significant results state a preference for making a choice over staying undecided. Bold p-values denote attraction by the pathogen side. Attractivity of the lures itself is tested on the right (no choice *vs*. choice). Significant results state a preference for making a choice over staying undecided.

| Life stage | Pathogen combination | Food – food & pathogen |  |  |  | No choice – choice |  |  |  |
| --- | --- | --- | --- | --- | --- | --- | --- | --- | --- |
| | | $\chi^2$ | df | p | N | $\chi^2$ | df | p | N |
| Adult females | *Beauveria* – *D. sulphureus* | 4.5 | 1 | **0.034** | 32 | 24.03 | 1 | < 0.001 | 35 |
|  | *Beauveria* – *R. canadensis* | 5.14 | 1 | **0.023** | 28 | 12.6 | 1 | < 0.001 | 35 |
|  | *Aspergillus* – *D. sulphureus* | 4.8 | 1 | 0.028 | 30 | 17.86 | 1 | < 0.001 | 35 |
|  | *Aspergillus* – *R. canadensis* | 0.36 | 1 | 0.549 | 25 | 6.43 | 1 | 0.011 | 35 |
|  | *Penicillium* – *D. sulphureus* | 24.5 | 1 | < 0.001 | 32 | 24.03 | 1 | < 0.001 | 35 |
|  | *Penicillium* – *R. canadensis* | 4.24 | 1 | 0.040 | 34 | 31.11 | 1 | < 0.001 | 35 |
| Larvae | *Beauveria* – *D. sulphureus* | 4.84 | 1 | 0.028 | 25 | 2.5 | 1 | 0.114 | 40 |
|  | *Aspergillus* – *D. sulphureus* | 2.13 | 1 | 0.144 | 30 | 10 | 1 | 0.002 | 40 |
|  | *Penicillium* – *D. sulphureus* | 6.53 | 1 | 0.011 | 30 | 10 | 1 | 0.002 | 40 |

**Figure 2:**
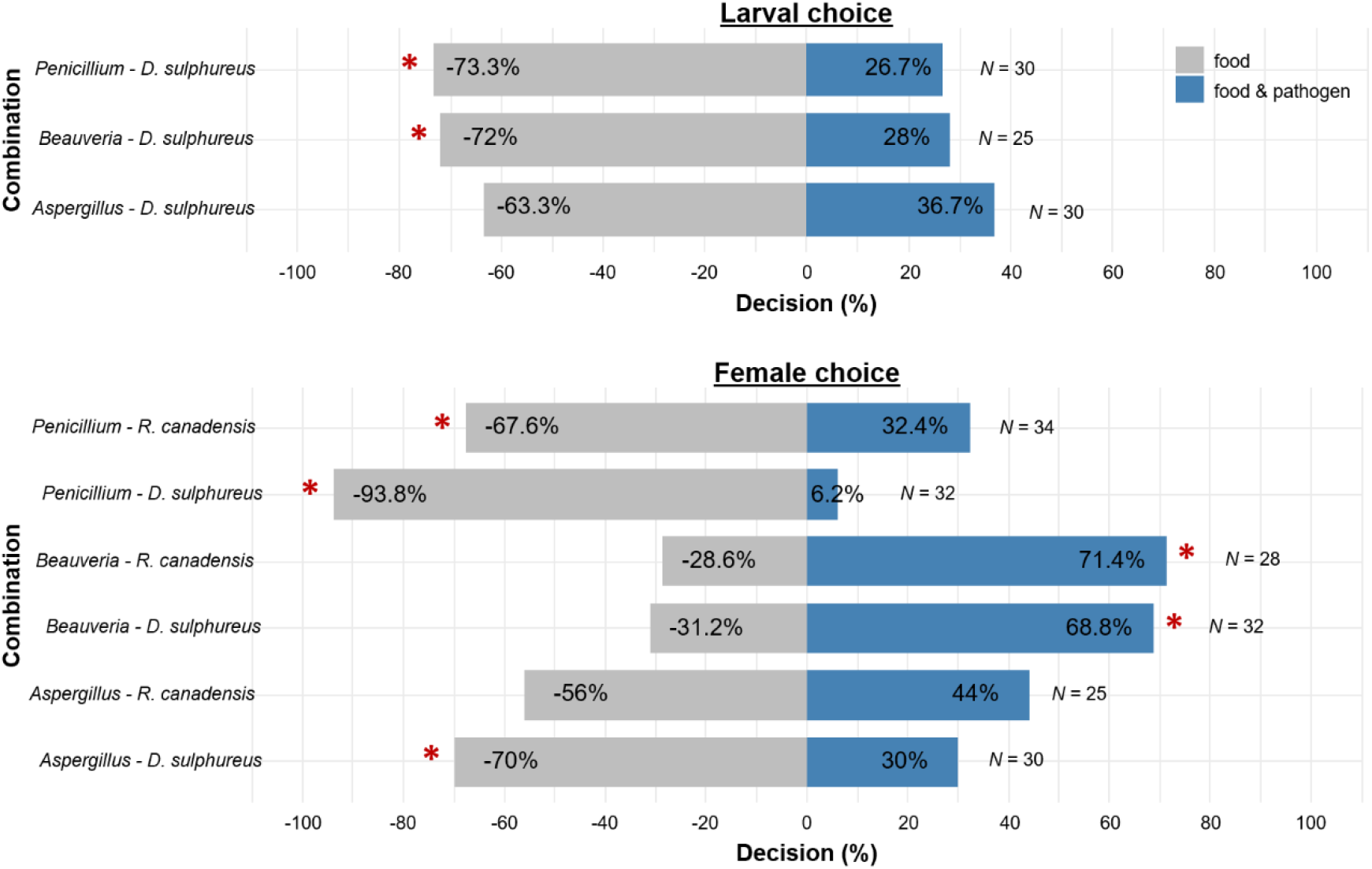
Decisions of larvae and adult females in the two-choice arenas. Percentage of larvae and adult females attracted to the positive control (= food fungus; left bars) or the repellence treatment (= food & pathogen; right bars) (*p < 0.05). N gives the total number of tested individuals. Non-deciding individuals were excluded here.

## Discussion

Overall, we were able to show that adult beetles and larvae of *X. saxesenii* responded positively to volatiles from their symbiotic fungi and even varied in their response to pathogens depending on their life stage. Larvae seem to be aware of their physical weakness compared to adults, especially in confrontation with the entomopathogen *Beauveria*. They do the opposite than their adult sisters and avoid contact with the entomopathogen. We suggest that female preference for *Beauveria* is a hygienic and social response to a serious hazard when present in the nest community. By confronting the entomopathogen, they have the chance to fight it before causing far-reaching damage. Various behaviours to reduce the establishment of parasites and pathogens like disinfection, relocation of dead and infected nestmates or waste management have already been shown in social insects (Cremer et al. 2007) and some of them are also present in *X. saxesenii* (Biedermann & Taborsky 2011; Nuotclá et al. 2019). Alternatively, it is possible that adult females are exploited by *Beauveria*, because it is a specialized entomopathogen that depends on infecting insects for survival. This is supported by the observation that volatiles of *B. bassiana* are apparently attractive also for other non-farming insects (Geedi et al. 2022 Nov 24).

Concerning the antagonistic fungi, both life stages were more attracted to the pure food fungus side, which can be interpreted in two ways. One would be a simple preference for a good nutritional food source, when having the choice, in particular when being away from the beetle community. This reaction might be different inside the nest. On the other hand, individuals could not see a severe hazard in the presence of the antagonists and both *P. commune* and *Aspergillus* sp. might be managed in other ways than active beetle treatment (e.g. passive application of antibiotic *Streptomyces griseus* (Grubbs et al. 2020)). Why adult individuals take no choice against *Aspergillus* when *R. canadensis* is used as a lure is unknown. It is possible that not volatiles of the pathogens themselves, but instead changed volatiles of the nutritional fungi when in presence with fungi are used as cues by the deciding beetles.

Future experiments may require a modified assay, which includes a more realistic in-nest situation with multiple conspecifics. Clearly, our results demonstrate a response of both larvae and adult ambrosia beetles towards other fungal volatiles. The decision to confront a potential threat or rather to avoid it may be based on more complex connections. Future studies should also investigate the volatiles of both fungal mutualists and pathogens and if and how co-occurrence influence these volatiles. This will also help to disentangle the single volatile compounds that are responsible for the attractions and repellences found.

## Acknowledgments

We thank our students Franziska Frass, Maximilian Philippi and Lara Quaas for the help during the experiments.

